# Functional phenomics and genetics of the root economics space in winter wheat using high-throughput phenotyping of respiration and architecture

**DOI:** 10.1101/2020.11.12.380238

**Authors:** Haichao Guo, Habtamu Ayalew, Anand Seethepalli, Kundan Dhakal, Marcus Griffiths, Xue-Feng Ma, Larry M. York

**Affiliations:** Noble Research Institute, LLC, 2510 Sam Noble Parkway, Ardmore, OK 73401

**Keywords:** GWAS, multi-trait, root architecture, root respiration, winter wheat

## Abstract

- The root economics space is a useful framework for plant ecology, but rarely considered for crop ecophysiology. In order to understand root trait integration in winter wheat, we combined functional phenomics with trait economic theory utilizing genetic variation, high-throughput phenotyping, and multivariate analyses.
- We phenotyped a diversity panel of 276 genotypes for root respiration and architectural traits using a novel high-throughput method for CO_2_ flux and the open-source software RhizoVision Explorer for analyzing scanned images.
- We uncovered substantial variation for specific root respiration (SRR) and specific root length (SRL), which were primary indicators of root metabolic and construction costs. Multiple linear regression estimated that lateral root tips had the greatest SRR, and the residuals of this model were used as a new trait. SRR was negatively correlated with plant mass. Network analysis using a Gaussian graphical model identified root weight, SRL, diameter, and SRR as hub traits. Univariate and multivariate genetic analyses identified genetic regions associated with aspects of the root economics space, with underlying gene candidates.
- Combining functional phenomics and root economics is a promising approach to understand crop ecophysiology. We identified root traits and genomic regions that could be harnessed to breed more efficient crops for sustainable agroecosystems.

## Introduction

Functional phenomics is an emerging transdisciplinary field that integrates physiology, high-throughput phenotyping, and computational biology in order to fill knowledge gaps about plant functioning (York, 2019). High-throughput phenotyping allows for large-scale data collection on plant form and function, and is often used for genetics within a species. Phenomics focuses on understanding variation in plant phenotypes, but often lacks analysis of the relation of phenotypes to function, even if quantitative genetics are employed. Therefore, functional phenomics is needed using statistical associations within high-dimensional phenomics data to infer how traits influence one another and physiological processes important for crop growth. Especially, root phenomics data and conceptual frameworks are lacking to understand their interactions and integration as described in York *et al.* (2013). The trait economics spectrum is a conceptual framework from ecology that could help explore trait integration in crops. In this context, economics refers to the balance among traits for resource acquisition and utilization, with an explicit treatment of the tradeoffs between pairs of traits (Reich, 2014). For example, in a controlled study of 74 plant species, a root economics spectrum was found in which root respiration correlated to percent nitrogen, root length per unit mass, and the decomposition rate of dried roots in soil (Roumet *et al.*, 2016). Recently, a root economics space was proposed formed by one axis representing whether to cooperate with fungal partners and a second representing the ‘fast’ or ‘slow’ strategies (Bergmann *et al.*, 2020). Interestingly, the first axis was partially driven by specific root length relating to construction cost, and the second axis by root nitrogen content, a proxy of specific root respiration and metabolic cost. Therefore, the root economics space is a useful framework for understanding carbon use efficiency in crop roots.

Roots are the interface between plants and soil, with a key function to extract nutrients and water that are required for plant productivity (Smith & De Smet, 2012; Meister *et al.*, 2014). However, there is a complex relationship between investing in the root system and plant productivity because roots have a cost. The fraction of newly fixed carbon from photosynthesis allocated to roots can exceed 50%, and this proportion to roots significantly increases under edaphic stress (Lambers *et al.*, 1996; Rachmilevitch *et al.*, 2015). Root system carbon costs can be classified as construction costs, including the structure of the roots and growth respiration, and maintenance costs, primarily respiration and exudation (Mooney, 1972; Sun *et al.*, 2020). For example, in wheat seedlings, 30% of net photosynthesis was measured as root construction and maintenance costs, such as respiration (Sawada, 1970). Therefore, optimizing construction and metabolism of the root system would have a significant impact on plant carbon use efficiency.

Specific root length is a measure of carbon expenditure to construct root length, often in units of m g^−1^. Specific root respiration standardizes respiration based on root length or mass, typically with units of nmol CO_2_ s^−1^ cm^−1^ or mg^−1^, respectively. Specific root length was found to have variation among a set of barley and wheat lines but the genetic contribution was not considered explicitly (Løes & Gahoonia, 2004), and was used for QTL analysis in common bean (Ochoa *et al.*, 2006). Across the plant kingdom, as much as 52% of photosynthates may be respired by plant roots during the same day, depending on species and environmental conditions (Lambers *et al.*, 1996). Plants respire photosynthetic substrates to produce carbon skeletons, usable energy, and chemical reduction needed for development (Amthor, 2000), which results in the consumption of oxygen and the release of carbon dioxide. A multicomponent framework has been suggested to divide respiration into (1) growth fraction, biosynthesis of new structural biomass and exudates, (2) maintenance fraction, translocation of photosynthate from sources to sinks, and cellular ion-gradient maintenance, (3) ion-uptake fraction, including uptake of ions, assimilation of N and S, and protein turnover (McCree, 1970; Thornley, 1970; Johnson, 1983; Poorter *et al.*, 1991; Amthor, 2000). As up to 60% of assimilated carbon is lost through respiration, strategies to minimize unnecessary respiratory activity could lead to substantial gains in crop productivity by enhancing plant carbon use efficiency (Amthor *et al.*, 2019; Weber & Bar-Even, 2019; Roell & Zurbriggen, 2020).

Variation in root respiration rates among crop species is due to the differences in root tissue density, anatomy, activity, chemistry, and structure (Ben-Noah & Friedman, 2018). Studies have shown that reducing root respiration through anatomical changes such as root cortical senescence of barley (*Hordeum vulgare* L.) and wheat (*Triticum aestivum* L.) (Schneider *et al.*, 2017) or reduction in root secondary growth of common bean (*Phaseolus vulgaris* L.) (Strock *et al.*, 2018) permit greater plant growth by improving phosphorus capture from low-phosphorus soils. Strategies have been proposed or used to reduce root respiratory carbon cost for improving plant performance, including making ion transport more efficient (Amthor *et al.*, 2019), manipulation of genes or enzymes involved in carbon metabolism in plant roots (Dorion *et al.*, 2017; Florez-Sarasa *et al.*, 2020), and using arbuscular mycorrhizal symbiosis to reduce root respiratory rate as well as increasing photosynthesis (Romero-Munar *et al.*, 2017). Root respiration that is not accounted for necessary plant functions might be referred to as luxury respiration.

Understanding the genetic bases of specific root length and respiration, among other traits, and their relationship to plant performance is of key importance for breeding more productive and resilient crop varieties to adapt to climate change. However, these traits have rarely been considered as a unit of phenotype for breeding or genetic mapping. Genome-wide association studies (GWAS) for respiratory traits will typically require many hundreds of plant variants, and measurement of respiratory traits at the same time of day and developmental stage (Scafaro *et al.*, 2017). Infrared gas analyzers for portable leaf photosynthesis or O_2_-electrodes techniques are commonly used to measure rates of root respiration (Poorter *et al.*, 1991; Strock *et al.*, 2018), but most of those protocols are low-throughput with costly instruments that have less flexibility for outputting convenient data formats. Addressing the need for rapid, cost-effective, large-scale root respiratory screening will require the development of both high-throughput root respiration measurement and data analysis capabilities, the combination of which will greatly strengthen functional phenomics by increasing statistical power and enabling genetic mapping (York, 2019).

Wheat, a member of the grass family, is an important cereal grown globally. Winter wheat in the Southern Great Plains of the United States is often grown as a dual-purpose crop for forage and grain production (Maulana *et al.*, 2019). Yield, protein content (Rajaram, 2001), disease resistance (Ellis *et al.*, 2014), and heat resistance (Maulana *et al.*, 2018) are major targets for modern wheat breeding and genetic improvement. Significant marker-trait associations for aboveground traits, such as yield and its components (Sukumaran *et al.*, 2018) and nitrogen use efficiency (Cormier *et al.*, 2016; Hawkesford & Griffiths, 2019), have been reported across the wheat genome. Indeed, considerable quantitative trait loci (QTL) associated with wheat root traits have been identified on nearly all chromosomes in variable environments (Hamada *et al.*, 2012; Bai *et al.*, 2013; Atkinson *et al.*, 2015; Maccaferri *et al.*, 2016; Xie *et al.*, 2017; Beyer *et al.*, 2019; Soriano & Alvaro, 2019). However, the genetic and functional basis of root traits still lag behind aboveground traits, and genetic variation of root construction and metabolic traits remains less explored. Accordingly, this study was conducted to (1) develop a high-throughput phenotyping platform that integrates a hydroponics growth system, infrared gas analyzers, custom gas chambers, a bead bath, flatbed scanners, analytical scales, and an R script for measuring specific root respiration, specific root length, and other root traits, (2) validate the platform using winter wheat to uncover heritable variation of root respiration and architectural traits, (3) emply functional phenomics to identify relations among traits and tissue-type dependencies, and (4) identify associated QTL/genes that drive root respiration and other root traits by performing GWAS.

## Materials and Methods

### Plant materials

The plant materials were selected from the hard winter wheat association mapping panel (HWWAMP) by the Triticeae Coordinated Agricultural Project (T-CAP). Two hundred seventy-six hard winter wheat cultivars and breeding lines were selected from the panel, which covers a broad range of selection and breeding history in the Great Plains of the USA.

### Experimental design

The 276 wheat lines were grown as two replicates in a single growth chamber with 552 plants, with the entire procedure repeated twice, for a total of four replicates and 1104 plants evaluated in this study. Each replicate was treated as a block for an overall experiment with a randomized complete block design. The two transplanting dates of seedlings into the growth hydroponics boxes were June 19 and October 4 in 2019. The details of the germination, growth, and sampling are given below.

### Growth conditions

Seeds were surface-sterilized in 0.5% NaOCl for 10 min and rinsed three times using deionized (DI) water, then pre-germinated in petri dishes with filter paper placed in darkness at 25°C for 3 d. Uniformly germinated seedlings were selected (Figure 1a), wrapped around the root-shoot junction with L800-D Identi-Plugs foam (Jaece Industries, NY, USA), plugged in a 15 mL conical centrifuge tube (VWR, Falcon®, catalog number: 21008-918) with the bottoms cut away from the “6 ml” mark, and transplanted to a hole cut into the lid of the growth system as described below (Figure 1b). A unique barcode label was affixed to each tube for sample identification. The hydroponics growth system consisted of a polypropylene divider box (inside dimensions: length of 38.10 cm, width of 22.86 cm, and height of 20.32 cm with a volume of 17.7 L) and a custom lid made from a PVC panel cut to fit in the top of a box (4.5 mm thick, by 250 mm wide, by 392 mm long with the corners cut off to accommodate the box’s rounded corners). Forty-eight holes with 18 mm diameter were drilled into the lid with a hole saw with equal spacing among holes. Twelve growth boxes were placed in a Conviron E-15 growth chamber (Conviron, Winnipeg, Canada) with a day:night cycle of 16:8 h, 25:20 °C, at a flux density at canopy level of ~400 μmol m^−2^ s^−1^. Each box was filled to the bottom of the lid with a nutrient solution containing (μM) 125 KH_2_PO_4_, 1125 KNO_3_, 500 CaCl_2_, 250 MgSO_4_, 11.5 H_3_BO_3_, 1.75 ZnSO_4_·7H_2_O, 2.25 MnCl_2_·4H_2_O, 0.08 CuSO_4_·5H_2_O, 0.03 (NH_4_)_6_Mo_7_O_24_·4H_2_O, and 19.25 Fe(III)-EDTA (C_10_H_12_N_2_NaFeO_8_). The nutrient solution was continuously aerated with an air pump attached to airstones submerged in each growth box, and the solution pH was maintained between 5.9 and 6.1 by additions of KOH or HCl throughout the experiment.

**Figure 1.**
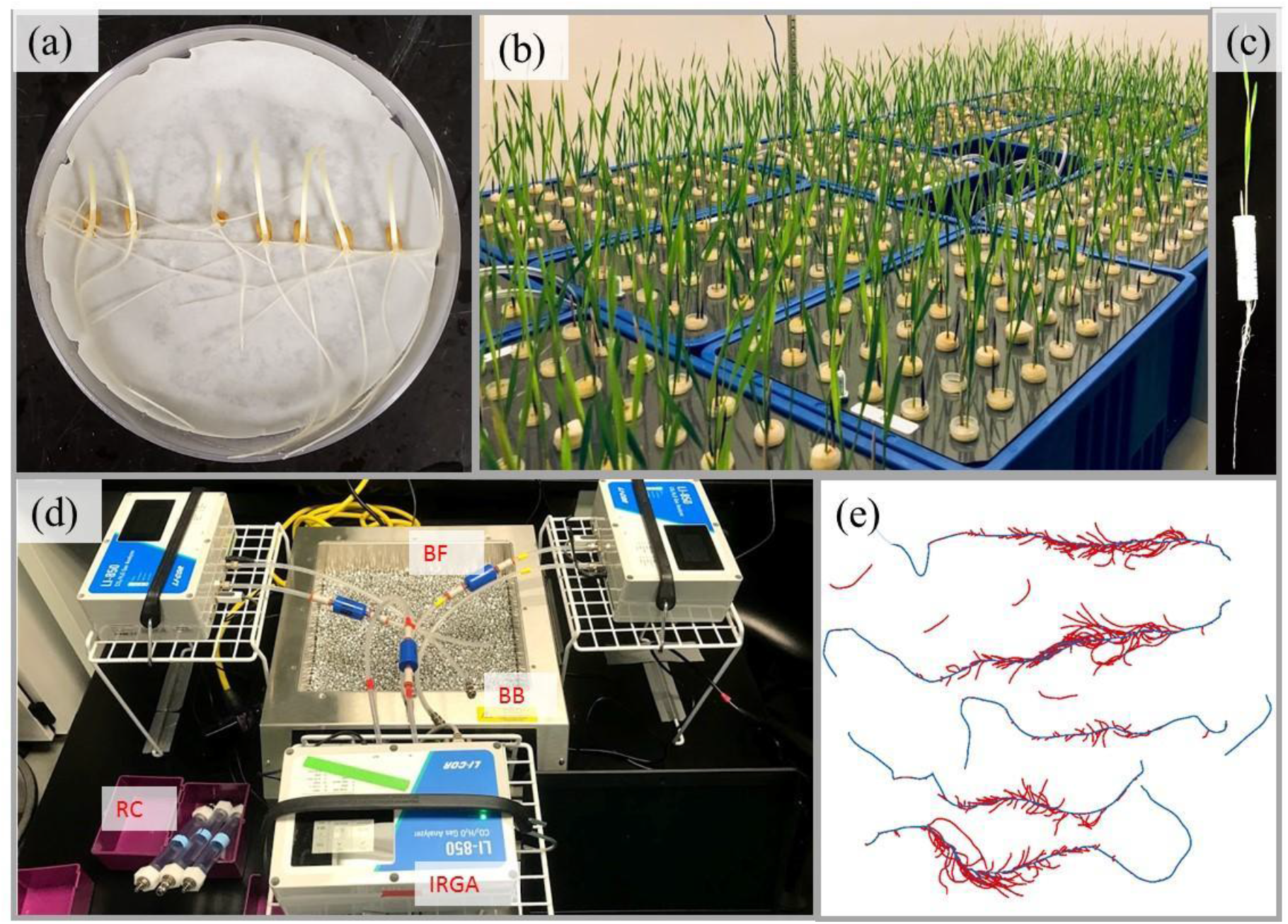
Platform for phenotyping root respiration and other root traits of wheat seedlings. (a) Wheat seeds were surface sterilized and pre-germinated in plate, (b) Seedlings were grown in aerated hydroponics for 10 days, (c) Shoot and roots of seedling 10 days after transplanting, (d) Root respiration was measured in an airtight chamber using a LI-850 with temperature control using a bead bath, (e) Distinguished axial roots (blue) from lateral roots (red) of scanned image using RhizoVision Explorer. IRGA: Infrared gas analyzer, RC: Root chamber, BB: Bead bath, BF: Balston Filter.

### High-throughput root respiration measurements

Ten days after transplanting (Figure 1c), plants were removed from the nutrient solution. Roots were immediately excised from shoots, blotted using tissue paper to remove excess water, placed in a 19 ml custom chamber, and then the chamber connected to an LI-850 CO_2_/H_2_O Analyzer (LI-COR Inc., NE, USA) (Figure 1d). The custom chamber was made from a 12.70 mm internal diameter clear polyvinyl chloride (PVC) pipe nipple (United States Plastic Corp., OH, USA) that was 152.4 mm in length with threaded ends. Holes were drilled into ½ inch FNPT nylon threaded caps (United States Plastic Corp., OH, USA) in order to accommodate insertion of quick-connect bulkhead male or female fittings (LI-COR Inc., NE, USA) with rubber grommets to create a seal. A Balston filter (LI-COR Inc., NE, USA) was inserted between the chamber and the analyzer to filter air. The chamber was buried in a Lab Armor bead bath (Model No 74300-714) filled with Lab Amor metallic beads with the temperature set at 28 °C. Beads were preferred to water in order to prevent contamination of the system with water. The chamber CO_2_ concentration was continuously recorded using LI-850 Windows software v 1.0.2 for 90 seconds at a rate of one reading per second. A USB barcode scanner (Taotronics, Fremont, CA) was connected to each laptop to acquire and save the datafile with the appropriate sample name encoded by the barcode affixed to the cut tube described above. Three infrared gas analyzers were used to allow simultaneous measurements in parallel to increase throughput.

In order to calculate the total respiration rate of a root sample from the individual text files containing the time series molar fraction of CO_2_, an R (R Core Team, 2018) script was developed in order to load each text file in a directory, do a series of computations, and output the total respiration rate. Total respiration rate (CO_2_ flux) was calculated using Equation 1.

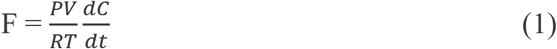

Where F is the CO_2_ flux in nmol s^−1^, P is the pressure in the chamber in kPa, V is the corrected chamber volume in milliliters, R is the ideal gas law constant in L kPa K^−1^ mol^−1^, T is air temperature in K, and dC/dt is the change in CO_2_ concentration on a molar basis with time (μmol mol^−1^ s^−1^). Chamber volume (V) was determined by subtracting the total root volume estimated using RhizoVision Explorer from the chamber volume. For root respiration analysis, the dead band (length of initial time to ignore) was set at 20 s. The slope of a linear regression fit to water-corrected CO_2_ concentration provided by the LI-850 analyzer over the corresponding observation time (20-90 s) using the *lm* function in R (R Core Team, 2018) is dC/dt. The protocol for the root respiration measurements and the R script for calculating total flux from a directory of text files are available at https://doi.org/10.5281/zenodo.4247873 (Guo *et al.*, 2020a).

After the root respiration measurements, roots from each plant were spread in a 5 mm layer of water in transparent acrylic trays and imaged with a flatbed scanner equipped with a transparency unit (Epson Expression 12000XL, Epson America) at a resolution of 600 dpi. Images were analyzed using RhizoVision Explorer version 2.0.2 (Seethepalli & York, 2020) with algorithms described by Seethepalli *et al.* (2020) with the options for image thresholding level, filter noisy components, threshold for root pruning being set at 205 pixel intensity, 0.2 mm^2^, and 1 pixel, respectively. A root diameter threshold of 0.30 mm was used to distinguish axial roots from lateral roots (Figure 1e).

Root traits extracted by the RhizoVision Explorer used in this study were number of root tips (Tip), number of branching points (BP), branching frequency (BF), total root length (TRL), axial root length (ARL), lateral root length (LRL), average diameter (AvgD), total root volume (TRV), axial root volume (ARV), lateral root volume (LRV), total root surface area (TSA), axial root surface area (ASA), and lateral root surface area (LSA). Branching frequency is determined by the software by dividing the number of branching points by total root length. Roots following scanning and shoots were dried at 60 °C for 3 days prior to dry weight determination. The oven-dried root mass and root length quantified using RhizoVision Explorer were used to calculate the specific root respiration (SRR) per unit of root dry mass (SRR_M; nmol g^−1^ s^−1^) and the specific root respiration per unit of root length (SRR_L; nmol m^−1^ s^−1^), respectively.

Root mass fraction (RMF) was calculated as root dry weight proportion of total plant dry weight. Specific root length (SRL) was calculated by dividing root length by the corresponding root dry weight. Lateral-to-axial root length ratio was calculated by dividing lateral root length by corresponding axial root length based on the diameter threshold provided during image analysis, and lateral-to-axial root volume ratio was calculated by dividing lateral root volume by corresponding axial root volume. Branching density (BD) was calculated by dividing root tips by axial root length. Root tissue density (RTD) was calculated by dividing root dry weight by root volume, which brought the total number of traits reported to 25 in this study.

Broad-sense heritability (*H^2^*) of each trait was calculated based on Falconer and Mackay (1996) as:

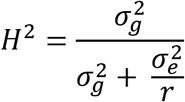

The variables 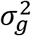, 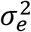 and *r*represent the variance of the genotype effect, variance of the local environment effect, and the number of replicates (blocks), respectively. The variances were obtained by fitting to a mixed model including genotype as a random effect and block as a fixed effect using the lme4 package (Bates *et al.*, 2014).

Principal component (PC) analysis and visualization of outputs were performed on the trait means of the 25 traits using the base function “prcomp” and the R package “factoextra” (Kassambara & Mundt, 2017). The first ten principal component scores were extracted for clustering and PC-based GWAS analysis (PC-GWAS).

### Network analysis

Due to highly correlated variables and singularities, root volume, surface area related traits, and lateral-to-axial root length ratio were dropped for network analysis. To assess the relationships among all the remaining 17 traits, we estimated pairwise Pearson’s correlation coefficients (*r*) of the traits and constructed a Gaussian graphical model for network analysis.

Network analysis with a Gaussian graphical model is more likely to capture causality and precursor/product relationships in trait networks relative to standard correlation analyses (Krumsiek *et al.*, 2011; Carlson *et al.*, 2019). The network analysis and the visualization of trait relationships were carried out with the R package ‘qgraph’ (Epskamp *et al.*, 2012). Outdegree is the number of connections that a trait node has to other trait nodes. Betweenness centrality quantifies the number of times a trait node acts as a bridge along the shortest path between two other trait nodes.

### Multiple linear regression analysis

Multiple linear regression analysis was employed to determine how total respiration can be partitioned into the contributions from root tissue types. The total axial root volume, lateral root axis volume (minus the tip volume), and lateral root tip volume were considered as the dependent variables while the total root respiration was the independent variable. The number of lateral root tips was estimated by subtracting 4 from the number of root tips supplied by RhizoVision Explorer, assuming that the typical wheat seedling had 4 seminal roots. This number of lateral roots was multiplied by 0.01 mm^3^ in order to assign a small volume to the lateral root tips, which were assumed to be more active based on previous research. Lateral root axis volume was determined by subtracting lateral root tip volume from the total lateral root volume. Based on visual evaluation of feature images in RhizoVision Explorer, total lateral root volume and total axial root volume were assumed as the volumes of the diameter ranges ≤ 0.3 mm or > 0.3 mm, respectively. The “stepAIC” function as implemented in R package “MASS” (Ripley *et al.*, 2013) was used for the stepwise regression and revealed this full model as being the most parsimonious, so residuals of this model were used as an additional trait (SRR_R) for subsequent analysis. SRR_R is the respiration that is not accounted for after considering root system architecture and root tissue dependency.

### SNP Genotyping

High-density single-nucleotide polymorphism (SNP) markers from the wheat 90K SNP genotyping array were obtained from Genotype Experiment “TCAP90K_HWWAMP” of The Triticeae Toolbox database (https://triticeaetoolbox.org/wheat/). Data constituting 21,555 SNPs were filtered to exclude markers with missing data greater than 50% and minor allele frequency less than 5%, resulting in 16,058 makers that were used in the association analysis. The map positions for the SNP markers used in this study were based on the consensus map developed using a combination of eight mapping populations (Wang *et al.*, 2014).

### Genome-Wide Association Analysis

Three genome-wide association analysis approaches were employed to identify genomic regions associated with various root traits. The linear mixed model (LMM) in GEMMA (Zhou & Stephens, 2012; Zhou & Stephens, 2014) was used to test for association between SNPs and traits. The population relatedness matrix was estimated using the centered relatedness algorithm within GEMMA, and was chosen as a covariate in the model. A Wald test was performed to determine *p*-values.

Single-trait (Univariate) association testing was run for each of the 25 traits using mean phenotypic values and PC-GWAS was conducted using each of the first 10 PCs. Multi-trait (Multivariate) GWAS was carried out to increase the power of the association tests and to detect polymorphisms with potentially pleiotropic effects of trait-associated loci using the multivariate linear mixed effect modeling capabilities of GEMMA. The 25 traits were grouped into six multi-trait combinations based on their genetic correlations, or their structural and functional relationships (McCormack *et al.*, 2017; Ben-Noah & Friedman, 2018). Root dry weight and shoot dry weight were combined to form a biomass-related multi-trait set (biomass). Total root respiration, root dry weight, root mass fraction, number of root tips, axial root length, and branching density were combined to form a root-respiration-related multi-trait set (root respiration) because these traits had functional relationships based on network analysis and provide a broader picture of root respiration. Axial root length, lateral root length, axial root volume, lateral root volume, axial root surface area, and lateral root surface area were combined to form a root-morphology-related multi-trait set (morphology). Branching point, branching frequency, and branching density were combined to form a root-topology-related multi-trait set (topology). Specific root length, root tissue density, and average root diameter were combined to form a root-construction-related multi-trait set (construction). Root mass fraction, lateral-to-axial root length ratio, and lateral-to-axial root volume ratio were combined to form an allocation-related multi-trait set (allocation). Multi-trait association was conducted with GEMMA using the multivariate version of the same model used for single-trait associations.

Outputs from GEMMA were used to generate Manhattan and Quantile-quantile (QQ) plots using the R package “qqman” (Turner, 2014). As mentioned in many wheat studies (Maulana *et al.*, 2018; Beyer *et al.*, 2019), determining a significance cutoff threshold is one of the biggest challenges for GWAS. Significant QTL were initially tested based on a false discovery rate of 0.05 following a stepwise procedure, which is very stringent (Müller *et al.*, 2011). So, an unadjusted significance level of −log_10_ P ≥ 3.5 was used for detecting SNPs that are significantly associated with the traits.

### Identification of candidate genes

The sequences of significant markers associated with phenotypic traits were downloaded from the The Triticeae Toolbox database (Wang *et al.*, 2014), and were BLAST searched against Phytozome’s version 2.2 of the wheat genome and identified candidate genes located ±250 kb proximal to each identified marker. Candidate genes of interest were selected based on the criteria of close proximity to the SNP, and possible involvement in the regulation of root development.

### Data and statistical code availability

All trait data, GEMMA output, and R analysis scripts necessary for doing the statistical analysis and plotting are available at https://doi.org/10.5281/zenodo.4247894 (Guo *et al.*, 2020b).

### Results

Variations of root respiratory and architectural traits

Shoot dry weight (SDW), RDW, TDW, TRR, SRL, L-to-A_L, ASA, L-to-A_V, PC2, PC3, PC4, and PC7 exhibited normal distribution. Near normal distributions were observed for other root traits (Fig. S1). The root traits with more than 5-fold variation between maximum and minimum values in the wheat population were SRR_L, TRL, LRL, LRV, LSA, and BP. 3.2-fold and 2.2-fold variations were also observed in SRR_M and SRL in the wheat population, respectively. Broad-sense heritabilities ranged from 0.25 to 0.57 for the 25 traits (Table 1). The respiration residual, SRR_R, of a multiple regression fit (Figure 2a) that accounts for respiration not explained by root system architecture, had a heritability of 0.44. The maximum heritability was observed for SDW (0.57). The root traits with heritabilities greater than 0.50 were SRL, BP, and AvgD. Many strong correlations were observed among traits. Total root respiration had correlation values greater than 0.50 with RDW and TDW. Interestingly, specific root respiratory traits (SRR_L and SRR_M) had significant negative correlations with shoot, root, and total dry weight (Figure 2b,Figure 3).

**Table 1.** Summary statistics and units for shoot dry weight, total dry weight, and the 24 root traits characterized in this study.

| Trait | Abbreviation | Unit | Mean | Min | Max | H<sup>2</sup> |
| --- | --- | --- | --- | --- | --- | --- |
| Shoot dry weight | SDW | g | 0.039 | 0.018 | 0.059 | 0.57 |
| Total dry weight | TDW | g | 0.053 | 0.024 | 0.080 | 0.51 |
| Root dry weight | RDW | g | 0.014 | 0.006 | 0.022 | 0.39 |
| Total root respiration | TRR | nmol CO<sub>2</sub> s<sup>-1</sup> | 0.54 | 0.23 | 0.91 | 0.42 |
| SRR per root length | SRR_L | nmol CO<sub>2</sub> s<sup>-1</sup> m<sup>-1</sup> | 0.14 | 0.04 | 0.34 | 0.48 |
| SRR per root mass | SRR_M | nmol CO<sub>2</sub> s<sup>-1</sup> g<sup>-1</sup> | 39.86 | 23.32 | 74.79 | 0.32 |
| SRR residual | SRR_R | nmol CO<sub>2</sub> s<sup>-1</sup> | -0.0039 | -0.3623 | 0.27 | 0.44 |
| Specific root length | SRL | m g<sup>-1</sup> | 299.7 | 182.21 | 398.36 | 0.55 |
| Root mass fraction | RMF | % | 26.48 | 19.05 | 36.17 | 0.43 |
| Total root length | TRL | mm | 4125.91 | 1315.65 | 7861.83 | 0.47 |
| Axial root length | ARL | mm | 1456.2 | 655.59 | 2537.42 | 0.48 |
| Lateral root length | LRL | mm | 2669.71 | 660.06 | 5494.75 | 0.48 |
| Lateral-to-axial root length ratio | L-to-A_L | mm mm<sup>-1</sup> | 1.82 | 0.83 | 2.66 | 0.48 |
| Total root volume | TRV | mm<sup>3</sup> | 329.49 | 135.78 | 610.24 | 0.45 |
| Axial root volume | ARV | mm<sup>3</sup> | 243.9 | 113.64 | 459.5 | 0.46 |
| Lateral root volume | LRV | mm<sup>3</sup> | 85.59 | 22.15 | 164.09 | 0.40 |
| Lateral-to-axial root volume ratio | L-to-A_V | mm<sup>3</sup> mm<sup>-3</sup> | 0.36 | 0.17 | 0.53 | 0.45 |
| Total root surface area | TSA | mm<sup>2</sup> | 3680.62 | 1348.64 | 6477.28 | 0.45 |
| Axial root surface area | ASA | mm<sup>2</sup> | 2034.03 | 934.42 | 3639.73 | 0.47 |
| Lateral root surface area | LSA | mm<sup>2</sup> | 1646.59 | 414.22 | 3266.23 | 0.44 |
| Average root diameter | AvgD | mm | 0.29 | 0.25 | 0.37 | 0.53 |
| Number of root tips | Tip | n | 399.61 | 161.67 | 710 | 0.40 |
| Number of branch points | BP | n | 931.66 | 283 | 1992.5 | 0.54 |
| Branching frequency | BF | n mm<sup>-1</sup> | 0.22 | 0.18 | 0.31 | 0.45 |
| Branching density | BD | n cm<sup>-1</sup> | 2.81 | 1.75 | 5.83 | 0.25 |
| Root tissue density | RTD | g cm<sup>-3</sup> | 0.04 | 0.03 | 0.06 | 0.30 |

H<sup>2</sup>, broad-sense heritability.

**Figure 2.**
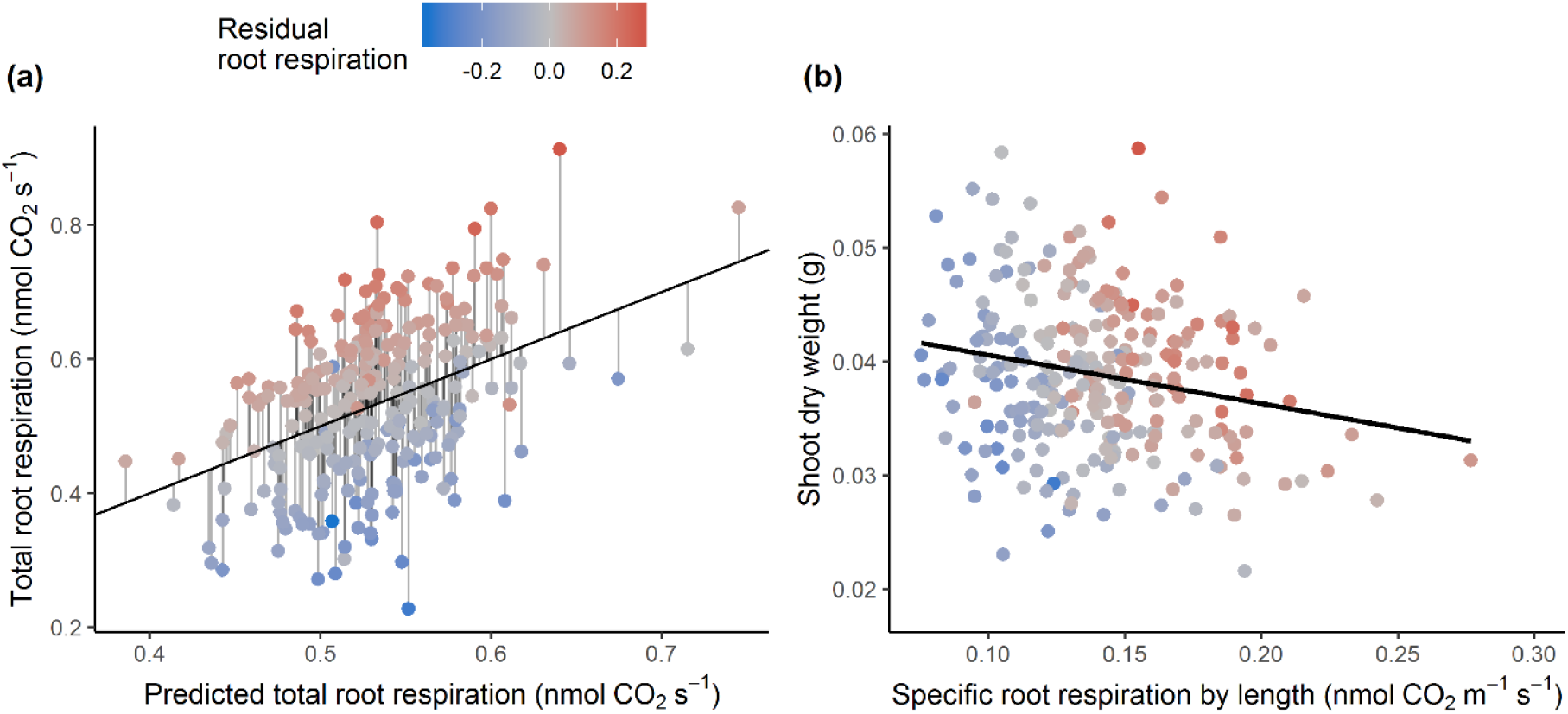
(a) The relationship between predicted total root respiration and total root respiration, and deviations from the relationship results in new trait SRR_R, (b) Regression between specific root respiration by length and shoot dry weight.

**Figure 3.**
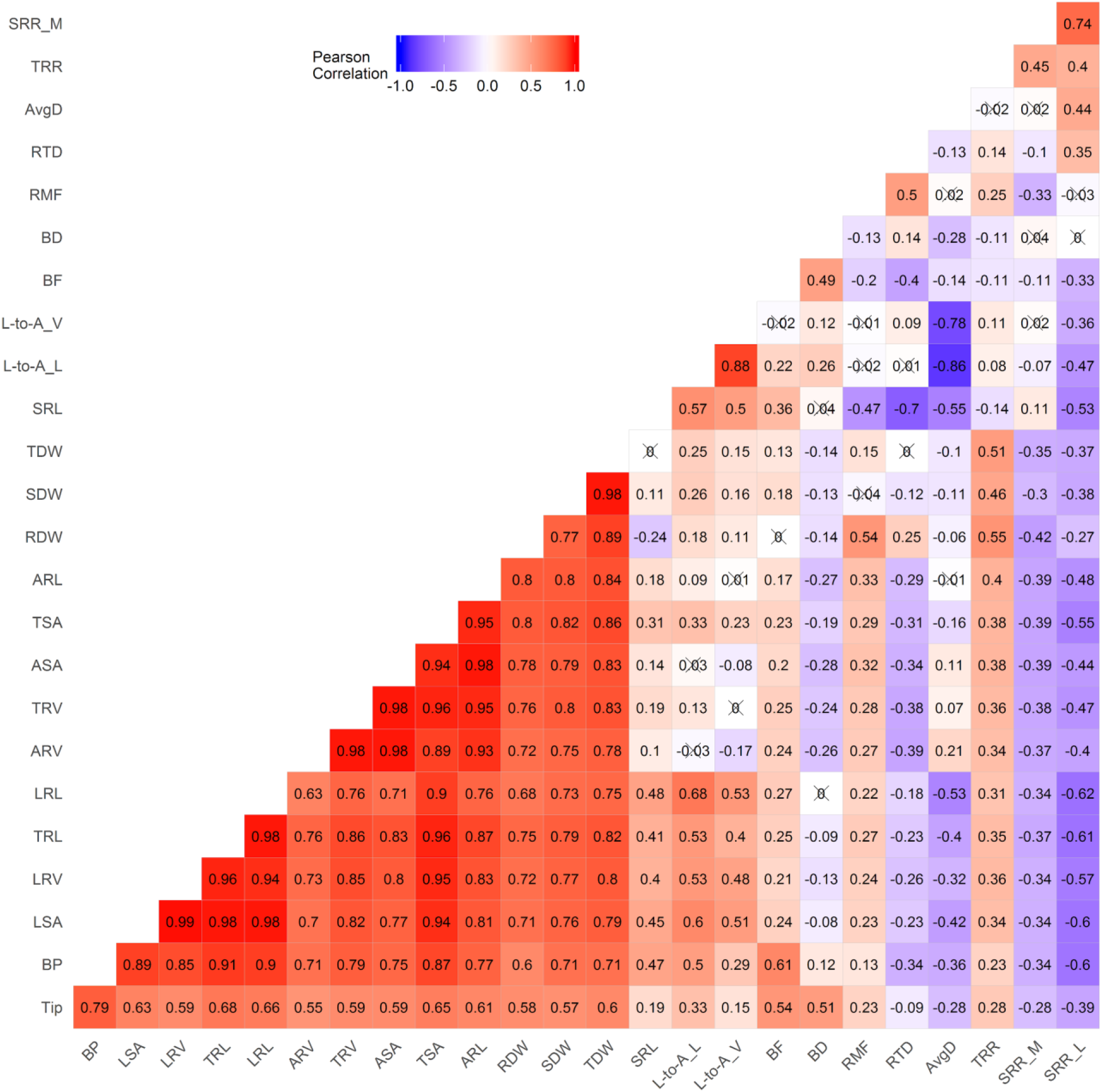
Pairwise Pearson correlation of selected traits of TCAP winter wheat seedlings. The number represents the correlation values. Value marked with symbol × means correlation is not significant at p = 0.05. Bright red to bright blue indicates highly positive to highly negative correlations, respectively. Trait abbreviations are as in Table 1.

Principal component analysis of the traits was conducted to further identify the major linear trait combinations that maximize the multivariate variation, and the first ten PCs collectively explained 98.8% of the total variance. PC1, PC2, PC3, and PC4 explained 49.9%, 17.5%, 9.3%, and 7.7% of the total variance, respectively (Figure 4a). Plant size-related traits including TSA, TRL, TRV, TDW, RDW, and SDW had important contributions (>5%) to PC1. In contrast, PC2 was largely driven by two construction cost related traits AvgD and SRL, which had contributions of 18% and 15%, respectively (Figure 4b). Traits with greater than 7% contributions to PC3 were the construction cost trait RTD (22%), three root respiration traits (TRR, SRR_L, and SRR_M), and branching trait BF (14%). PC4 was predominantly driven by SRR_M and SRR_L that represent metabolic costs, which had contributions of 24% and 14% to the component, respectively (Figure 4c).

**Figure 4.**
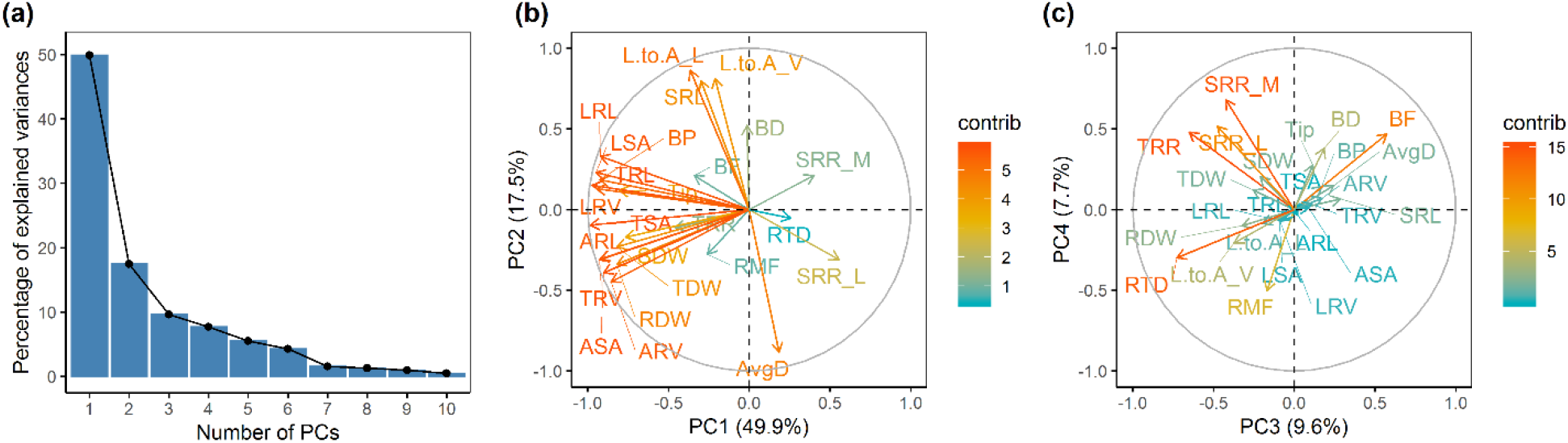
(a) Scree graph showing the percentage of variance explained by each of the first 10 principal components, PCA variable contribution plots showing the (b) first and second PCs and (c) third and fourth PCs, where vectors indicate relative weightings of the variables. Trait abbreviations are as in Table 1.

### Multiple linear regression partitions respiration among root tissue types

Multiple linear regression analysis was employed to determine the respective contributions of lateral root tip, lateral root axis, and toal axial root volumes to total root respiration, and to provide the SRR_R trait. The resulting model (*p* < 2.2e-16) explains 14.5% of the variation in total root respiration. Axial root volume, lateral root volume, and lateral root tip volume were all significant explanatory variables (*p* = 0.001, 1.37e-05, and 0.033, respectively). The average specific root respiration rate on a volume basis of lateral root tips was 30.5 and 8.1 times the rates of axial roots and lateral roots, respectively, as determined from comparing slopes in the model. The residuals represent respiration not accounted for by average tissue dependencies within the diversity panel, which we hypothesized to have a genetic component.

### Trait correlation network

In addition to the correlation analyses, a network analysis based on a Gaussian graphical model was performed to account for the conditional dependencies between the investigated traits. The traits exhibiting outdegree greater than 2.0 were AvgD, RTD, ARL, SRR_M, and SRL in descending order (Table S1). Average root diameter showed the highest betweenness, connecting a root branching subnetwork via ARL, and a biomass subnetwork via RMF. SRL also exhibited a high betweenness, by connecting other groups of traits belonging to root respiration, biomass, root morphology, and topology. Consistent with Pearson correlation analysis, SRR_M was weakly connected with root dry weight, total dry weight, and RMF. SRL was negatively and positively correlated with SRR_L and SRR_M, respectively (Figure 5). In contrast to the Pearson correlation analysis (Figure 3), no direct network connection was observed between shoot dry weight and root respiratory and architectural traits (Figure 5).

**Figure 5.**
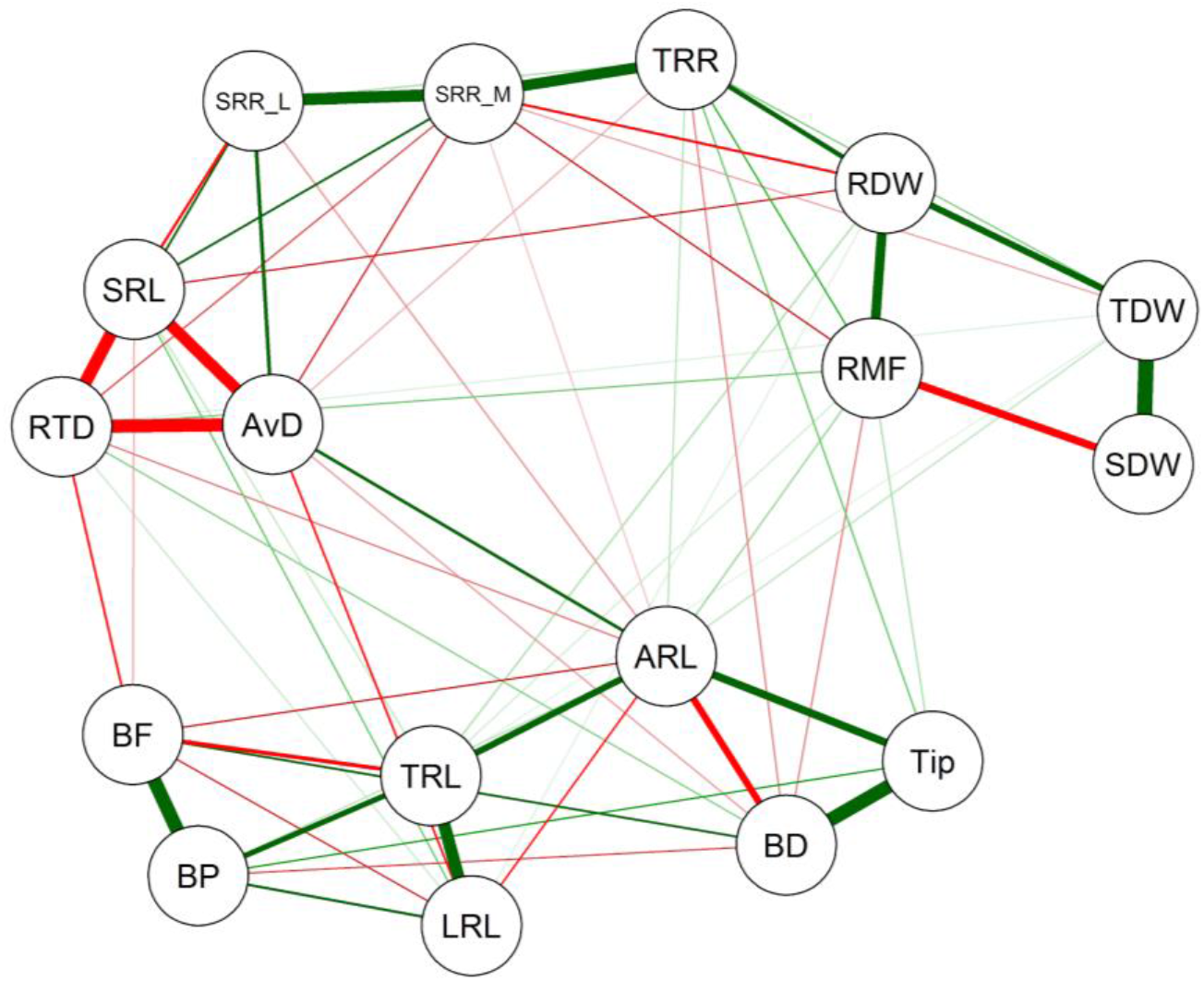
Trait correlation network constructed from Gaussian graphical model. Red and green edges show negative and positive correlations, respectively. Cutoff was set at 0.15. Trait abbreviations are as in Table 1.

### Genome-wide association analysis

Multi-trait GWAS on the six sets of traits identified 140 SNPs while the single-trait GWAS of 25 traits identified 234 significantly associated SNPs (−log_10_ P = 3.5). GWAS based on the first 10 PCs identified 79 SNPs that passed the −log_10_P of 3.5, and the majority of these detected SNPs were associated with PC1, PC2 or PC9 (Figure 6a,Table S2). Sixty-nine percent of the significantly associated SNPs in multi-trait approach and 56% of the SNPs in PC-GWAS were represented in the single-trait GWAS (Figure 6). Overall, multi-trait GWAS and PC-GWAS identified 77 additional, unique SNPs that were not uncovered by the 25 univariate analyses (Figure 6a,Figure S2).

**Figure 6.**
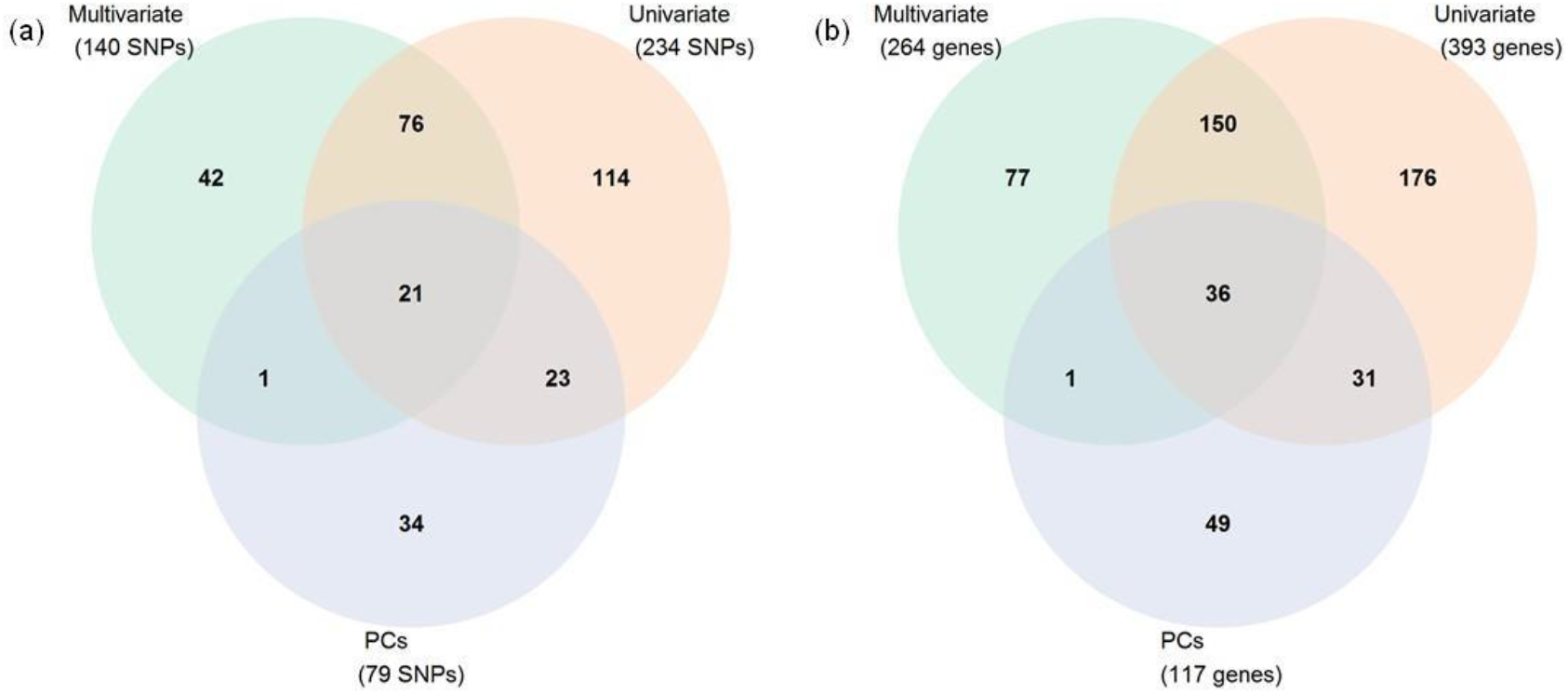
Venn diagram of (a) associated SNPs cutoff set at −log_10_P=3.5, (b) genes identified using cutoff set at −log_10_P= 3.5 comparing significantly univariate of mean 25 traits, univariate analyses of 10 principal components (PCs), and multivariate of 6 trait combinations.

Four significant markers associated with SRR_M were identified on chromosomes 1B, 4B and 4D (Figure 7a). There were no genes underlying the top two largest −log_10_P signals on chromosomes 1B and 4B, while the third largest −log_10_P signal (IWA430) on chromosome 4D was encoding for four underlying proteolysis genes involved in cellular protein catabolic process (Table S3). Seven significant markers associated with SRR_L were identified on chromosomes 4B and 5A. The marker (Excalibur_c100336_106) with the largest −log_10_P signal on chromosome 4B, which co-associated with SRR_M, had no known underlying gene. Six genes underlying the next two largest −log_10_P signals on chromosome 5A were annotated with functions as ATP binding, protein binding, and protein kinase activity (Table S3). Three additional significant markers associated with SRR_R were detected on chromosomes 1A and 1B (Table 2). Three genes underlying the largest −log_10_P signal (Kukri_c10453_875) on chromosome 1A were associated with processes of DNA transcription regulation (Table S3). There were no genes underlying the other two markers. Multi-trait GWAS for root respiration identified 20 additional markers on chromosomes 1A, 1B, 2B, 3D, 4A, 4B, 5B, 6A, and 7A (Figure 7a). There were no known genes underlying the largest −log_10_P signal Excalibur_c5139_198 on chromosome 1A, and four genes underlying the following two largest −log_10_P signals on chromosomes 1A and 1B were annotated with functions as protein kinase activity and ADP binding (Table S3).

**Table 2.** Subset of significant SNP markers identified from multi-trait GWAS and univariate GWAS of single-trait by selecting top three SNPs of each trait.

| Trait | Model | Markers | Chr | MAF | p value |
| --- | --- | --- | --- | --- | --- |
| SRR_M | Univariate | Excalibur_c100336_106 | 4B | 0.110 | 1.91E-05 |
| SRR_M | Univariate | IAAV5776 | 1B | 0.056 | 9.95E-05 |
| SRR_M | Univariate | IWA430 | 4D | 0.438 | 2.11E-04 |
| SRR_L | Univariate | Excalibur_c100336_106 | 4B | 0.110 | 1.70E-05 |
| SRR_L | Univariate | CAP12_c956_61 | 5A | 0.112 | 2.04E-04 |
| SRR_L | Univariate | BS00066434_51 | 5A | 0.146 | 2.19E-04 |
| SRR_R | Univariate | Kukri_c10453_875 | 1A | 0.281 | 7.51E-05 |
| SRR_R | Univariate | IWA6965 | 1B | 0.064 | 9.86E-05 |
| SRR_R | Univariate | RAC875_c42206_305 | 1B | 0.064 | 9.86E-05 |
| Respiration | Multivariate | Excalibur_c5139_198 | 1A | 0.213 | 5.69E-06 |
| Respiration | Multivariate | tplb0048b10_1365 | 1B | 0.064 | 4.35E-05 |
| Respiration | Multivariate | Ex_c4876_1221 | 1A | 0.248 | 9.36E-05 |
| SRL | Univariate | RAC875_c63889_486 | 7A | 0.202 | 7.58E-05 |
| SRL | Univariate | GENE-1220_457 | 2A | 0.064 | 1.29E-04 |
| SRL | Univariate | RFL_Contig5917_2369 | 2A | 0.071 | 1.73E-04 |
| AvgD | Univariate | IWA7907 | 7B | 0.190 | 1.95E-04 |
| AvgD | Univariate | IWA4438 | 7A | 0.083 | 1.97E-04 |
| AvgD | Univariate | Tdurum_contig61864_1352 | 7A | 0.082 | 2.03E-04 |
| RTD | Univariate | GENE-0249_161 | 1B | 0.272 | 6.84E-05 |
| RTD | Univariate | IWA614 | 7A | 0.277 | 1.97E-04 |
| RTD | Univariate | Kukri_c20062_389 | 1D | 0.165 | 2.06E-04 |
| Construction | Multivariate | BS00082644_51 | 3B | 0.247 | 4.91E-05 |
| Construction | Multivariate | GENE-0249_161 | 1B | 0.272 | 1.65E-04 |
| Construction | Multivariate | IWA6076 | 2B | 0.273 | 1.72E-04 |
| BF | Univariate | CAP7_c1083_283 | 1A | 0.140 | 1.74E-05 |
| BF | Univariate | Kukri_c29121_226 | 1A | 0.140 | 1.98E-05 |
| BF | Univariate | Kukri_c53935_265 | 1A | 0.136 | 3.47E-05 |
| BP | Univariate | IWA1464 | 1D | 0.147 | 6.00E-07 |
| BP | Univariate | BS00032149_51 | 1D | 0.133 | 7.39E-07 |
| BP | Univariate | IWA2164 | 1D | 0.150 | 1.31E-06 |
| BD | Univariate | Tdurum_contig49608_1185 | 4B | 0.143 | 1.59E-04 |
| BD | Univariate | BS00063973_51 | 5A | 0.404 | 2.63E-04 |
| BD | Univariate | Excalibur_c33173_557 | 2D | 0.205 | 2.91E-04 |
| Topology | Multivariate | BS00032149_51 | 1D | 0.133 | 6.15E-06 |
| Topology | Multivariate | IWA1464 | 1D | 0.147 | 6.95E-06 |
| Topology | Multivariate | IWA2164 | 1D | 0.150 | 1.74E-05 |

Chr, Chromosome; MAF, minor allele frequency.

**Figure 7.**
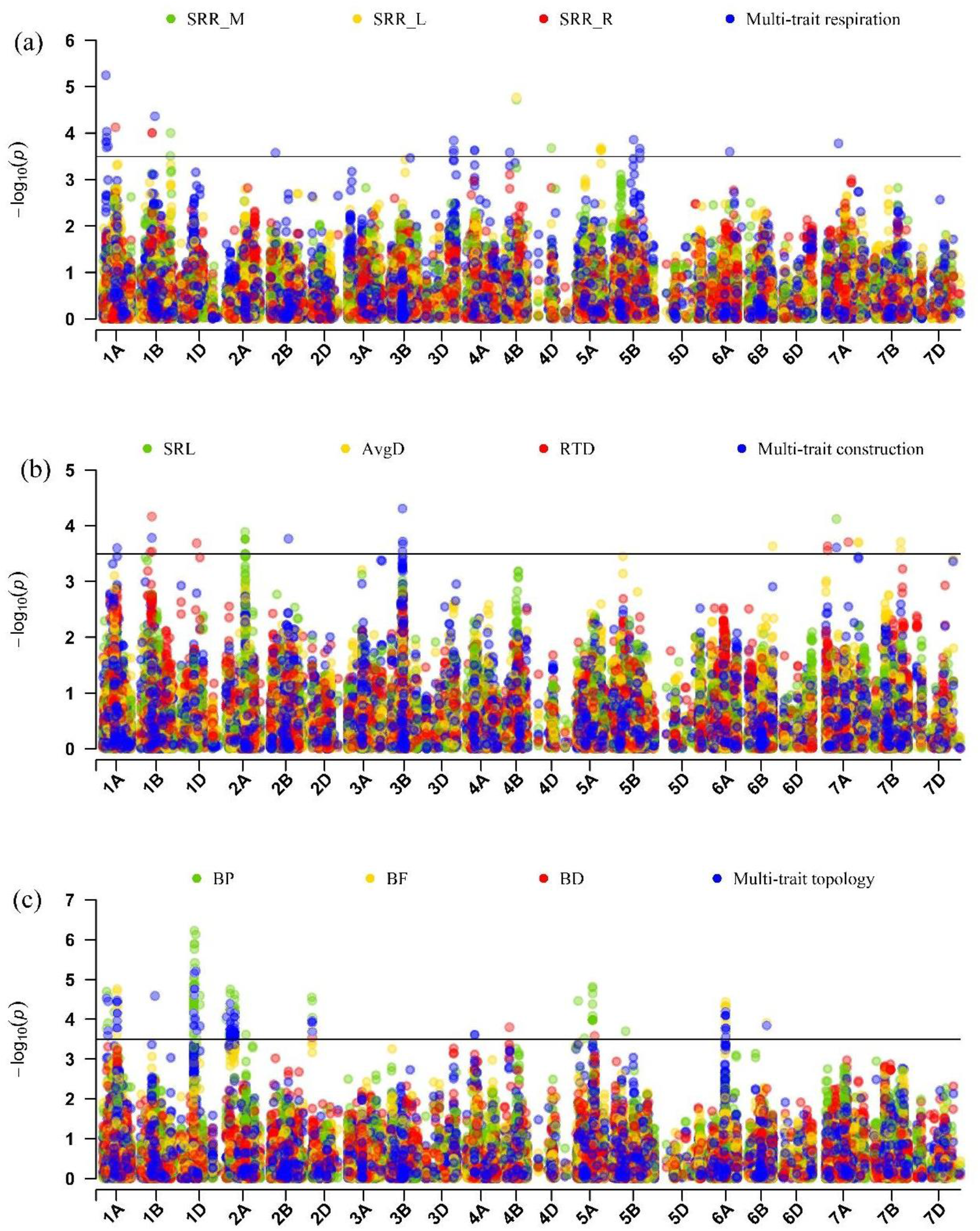
Manhattan plot of GWAS conducted on traits (a) SRR by mass, SRR by length, residuals of total respiration vs volume of different segments, and multi-trait combination for root respiration, (b) SRL, AvgD, RTD, and multi-trait combination for root construction, (c) BP, BF, BD, and multi-trait combination for root topology of TCAP wheat population. Each dot represents a SNP. The horizontal black line indicates the threshold of significance at − log_10_P = 3.5. Trait abbreviations are as in Table 1.

Ten significant markers associated with single-trait SRL were identified on chromosomes 2A (9 markers) and 7A, and seventeen genes underlying the top three largest −log_10_P signals on chromosome 2A and 7A have functions related to protein binding, calcium ion binding, polysaccharide binding, and ATP binding (Figure 7b,Table S3). Five significant markers associated with single-trait AvgD were identified on chromosomes 6B, 7A, and 7B. Only one of the top three largest −log_10_P signals on chromosome 7Ahad three underlying genes, which were annotated with function as protein binding. Seven significant markers associated with single-trait RTD were identified on chromosomes 1B, 1D, and 7A, and eight genes underlying the top three largest −log_10_P signals on chromosomes 1B, 1D, and 7A were annotated as zinc finger CW-type coiled-coil domain protein and integral membrane Yip1 family protein. Multi-trait GWAS for root construction identified eight markers on chromosomes 1A, 1B, 2B, 3B, and 7A, with one marker (GENE-0249_161) on 1B co-associated with single-trait RTD, and another marker (RAC875_c63889_486) on 7A co-associated with single-trait SRL (Table 2). Eight genes underlying the top three largest −log_10_P signals on chromosomes 1B, 2B, and 3B were annotated as regulators of VPS4 activity protein-related and potassium ion transmembrane transport (Table S3).

Thirty-four significant markers associated with single-trait BF were identified on chromosomes 1A, 2A, 2D, 6A, and 6D, and seven genes underlying the top two largest −log_10_P signals on chromosome 1A involved in biological processes such as oxidation-reduction, steroid biosynthetic process, and DNA-binding process. We detected 134 markers for single-trait BP on chromosomes 1A, 1D, 2A, 2D, 5A, 5B, and the top three largest −log_10_P signals and underlying nine genes were all observed on chromosome 1D, which also co-associated with multi-trait root topology. Three significant marker associations were detected for single-trait branching density on chromosomes 2D, 4B, and 5A (Figure 7c), and five genes underlying the three markers had annotations indicating involvement in oxidation-reduction biological processes. Multi-trait GWAS for root topology identified 84 significant markers, with 80 markers co-associated with single-trait BF or BP (Figure 7c). Significant marker associations and underlying genes were also detected for multi-trait biomass, multi-trait allocation, multi-trait morphology, for all PC-traits except PC8, and for the other single traits (Figure 6,Figure S2, Table S3).

## Discussion

Reducing the metabolic and construction carbon costs of roots has become a realistic engineering strategy for crop breeding to increase yield and promote plant growth (Lynch, 2013; Lynch, 2018; Amthor *et al.*, 2019). However, the genetic and functional basis of root respiration traits still lags behind architectural root traits. Scaling up the throughput of root respiration phenotyping will strengthen functional phenomics greatly by increasing statistical power and enabling genetic mapping (York, 2019). The platform we developed facilitates high-throughput phenotyping of root respiration, with integration of cost-effective equipment and an R script for data processing, and allowed throughput of about 25 samples person^−1^ hour^−1^. The use of a bead bath for controlling temperature avoids the risk found when using a water bath of water entering the respiration chamber and contaminating the gas analyzer. We observed 8.5-fold variation for SRR_L and 3.2-fold variation for SRR_M in the wheat panel. In previous work, root respiration was measured mostly using single root segments (Poorter *et al.*, 1991; Strock *et al.*, 2018), and there was little information about how different root types impact respiration of whole root systems. Considering the difficulty of separating different root tissue segments from whole root systems for maintaining high-throughput, multiple linear regression was used to predict the contributions of root tissue types to total root respiration of wheat seedlings on average within the panel. We found that the lateral root tips had much greater respiration than axial root tissue or lateral root axis tissue(≤ 0.3 mm), which supports findings in woody plants that root tip meristems consume about 15 times more O_2_ than the rest of the root system (Mancuso & Boselli, 2002; Aguilar *et al.*, 2003; Burton *et al.*, 2012).

Correlation network analyses have been widely used in biology and social sciences to capture causality and precursor/product relationship patterns in functional traits. Despite the elegance of this approach, only a few studies applied network theory to plant root traits (Poorter *et al.*, 2014; Messier *et al.*, 2017; Carlson *et al.*, 2019; Kleyer *et al.*, 2019). In addition to root dry weight, SRL, and average diameter, SRR_M, which is rarely used in functional trait analysis, was identified as one of the hub traits and had substantial effects on the plant phenotype as a whole. Consistent with previous work, SRL correlated with root dry weight, root diameter, branching, and root tissue density (Reich, 2014; Kramer-Walter *et al.*, 2016). In addition, we found that SRL also can be an indicator of root respiration on either a mass or length basis. Shoot biomass only had a strong positive correlation with total biomass and a negative correlation with root mass fraction in the network, which may indicate that the formation of wheat seedling shoot biomass was mostly independent, and also indicates that reducing or otherwise optimizing the allocation of resources to the root could be a strategy to improve shoot growth (Guo & York, 2019). Counterintuitively, driving shoot growth with such a strategy may actually maintain root mass and total metabolic burden, or even increase these total costs, but with less proportion relative to the shoot. This framework of carbon use efficiency represents an untapped positive feedback loop for plant growth. Interestingly, network, principal component, and regression analyses all showed that SRR_M was negatively correlated with total dry weight, suggesting that reducing respiratory carbon could potentially increase whole-plant growth (Lynch, 2015; Amthor *et al.*, 2019).

Multi-trait GWAS has recently gained more attention because it often boosts the power to detect SNPs and assesses the full spectrum of traits that are affected by trait-associated variants (Porter & O’Reilly, 2017), which can be particularly useful for challenging physiological traits (Chhetri *et al.*, 2019). Combining traits related to respiration, multi-trait association analysis identified 20 unique significant associations while single-trait GWAS detected 13 unique significant associations for all SRR traits. The findings potentially reveal the pleiotropic effects of genes near significantly associated SNPs on root respiration. The marker tplb0048b10_1365, the second-largest −log_10_P signal associated with multi-trait root respiration, was reported to be associated with nitrogen deficiency tolerance in wheat seedlings (Ren *et al.*, 2018). Multiple annotated genes underlying significant SRR_L and SRR_M associated SNPs are annotated with functions in protein catabolism, protein binding, ADP, and ATP binding, which are related to cellular respiration (Araújo *et al.*, 2011), root meristem activity (Xu *et al.*, 2017) or root senescence (Liu *et al.*, 2019).

GWAS for root architectural traits have gained increasing attention in wheat, and several QTL/genes in wheat have been found to associate with root architectural and morphological traits such as root length, root number, and root diameter across the genome (Maccaferri *et al.*, 2016; Ayalew *et al.*, 2018; Beyer *et al.*, 2019). Specific root length (SRL), AvgD, and RTD are important components of the root economic spectrum because they potentially provide information about root morphology and construction costs (Kramer-Walter *et al.*, 2016; McCormack *et al.*, 2017). Multiple genes underlying associated significant SNPs were identified as zinc finger protein, cytochrome p450 family member, and haloacid dehalogenase-like hydrolase family protein, which all play important roles in controlling wheat root growth and development (Kulkarni *et al.*, 2017; Li & Wei, 2020). Multiple genes underlying two markers (Kukri_c24648_262 and Kukri_c5113_1082), which were co-associated with TRL, LRL, TRV, LRV, TSA, LSA, BP, PC1, and multi-trait allocation and topology (Table S3), were annotated as a nucleoporin autopeptidase domain containing protein. Those genes may play distinct roles in nuclear transport and root elongation (Parry, 2014).

Root branching is a necessary developmental process for increasing the number of growing tips and defining the distribution of their meristem sizes (Pagès, 2014), with a large metabolic cost. Root branching was critical for plant survival and performance under abiotic conditions (Schneider *et al.*, 2020). Two genes (Traes_1AL_9CC946A58 and Traes_1DL_7EF27C52F) underlying the largest −log_10_P signal of BF were annotated as being involved in steroid biosynthesis, which may play a role in interacting with auxin signaling to promote lateral root growth (Vriet *et al.*, 2012; Wang *et al.*, 2018). Four genes underlying the marker BS00013534_51, which was co-associated with BF, BP, and multi-trait topology, were annotated as encoding protein kinase activity in wheat and threonine-protein kinase receptor precursor in rice. Interestingly, different genes with similar functions were found playing fundamental roles in lateral root formation and development (Atkinson *et al.*, 2014; Yu *et al.*, 2016; Pan *et al.*, 2020). Three candidate genes underlying single-trait branching frequency and multi-trait topology co-associated SNP (tplb0025i05_1836) were annotated as being involved in the activity of Rho guanine nucleotide exchange factors. Rho family members are well known as regulators of extracellular stimulus-dependent signaling pathways that affect gene expression, cell proliferation, actin cytoskeleton, cell cycle progression, and cell polarity (Berken & Wittinghofer, 2008).

A recent review outlined the emerging possibilities for targeted reducing unnecessary carbon loss for increasing yields (Amthor *et al.*, 2019), which was further supported by new simulation results indicating that substantial gains could be made by targeting plant respiration (Holland *et al.*, 2019). Therefore, an optimal root system will conform to economic cost-benefit analysis where the cost increment of allocation to the root system equals the benefit increment, measured as nutrient and water capture, or marginal photosynthesis (Bloom *et al.*, 1985). Recent work from the RIPE project has also shown it’s possible to increase photosynthesis by reducing photorespiration (South *et al.*, 2019) and increasing photosynthetic induction (Acevedo-Siaca *et al.*, 2020). We propose that combining strategies that increase photosynthesis and decrease ‘luxury’ root respiration could have synergistic and compounding influences on plant growth. The root economics space discussed above provides a useful framework for this strategy.

## Conclusions

We developed a high-throughput platform for measuring multiple traits within the root economics space, including root respiration and specific root length which are aspects of root metabolic and construction costs, respectively. Substantial, heritable variation exists within wheat, providing further evidence for intraspecific economics spectra. Employing the functional phenomics approach allowed leveraging genetic and phenotypic diversity to infer the increased contribution of lateral root tips to respiration, the negative relation of SRR to seedling mass, and network analysis that identified hub traits. Genome-wide association studies for the univariate traits uncovered several underlying genetic regions, while multivariate and PCA-based GWAS provided increased power to detect genetics of the root economics space itself for the first time to our knowledge. The SNPs associated with the traits may be useful for marker-assisted breeding. Candidate genes underlying significant SNPs associated with root respiratory, construction, and topology traits will require further research for reducing respiratory carbon loss and construction costs. We provide evidence that combining functional phenomics methods and trait economic theory has substantial potential to advance plant biology and crop breeding.

## Acknowledgements

This research was supported by the Noble Research Institute, LLC and the Samuel Roberts Noble Foundation. The authors wish to acknowledge David McSweeney and Karen Hartman of Greenhouse Core Facility for assistance provided during the experiment, Nick Krom of Scientific Computing Department for assistance with BLAST searches for candidate genes, and contributions from Yaxin Ge, Michael Cloyde, Na Ding, Xinji Zhang, Guangming Li, and Wangqi Huang for data acquisition and sampling.

## Author contributions

HG and LMY designed the experiments. X-FM provided the germplasm and expertise for genetic analysis. HG, AS, KD, and LMY conducted experiments. HG, MG, AS, KD, and LMY developed the respiration measurement protocol and R script for root respiration analysis. HG, AS, KD, HA, and LMY analyzed the experimental data. HG and LMY wrote the first draft of the manuscript, all authors made revisions, and all approved the final version.

## Supporting Information

**Figure S1** Histograms for the frequency distribution of 26 traits and 10 PC scores

**Figure S2** Manhattan plots of GWAS conducted on all traits

**Figure S3** Quantile-quantile (Q-Q) plots for all traits

**Table S1** Centrality measures of 17 traits from Gaussian graphical model

**Table S2** List of SNPs using a cutoff value set at −log_10_P=3.5

**Table S3** List of nearest genes underlying SNPs using a cutoff value set at −log_10_P=3.5

